## Supplemental Data for "KDM6B promotes activation of the oncogenic CDK4/6-pRB-E2F pathway by maintaining enhancer activity in MYCN-amplified neuroblastoma"

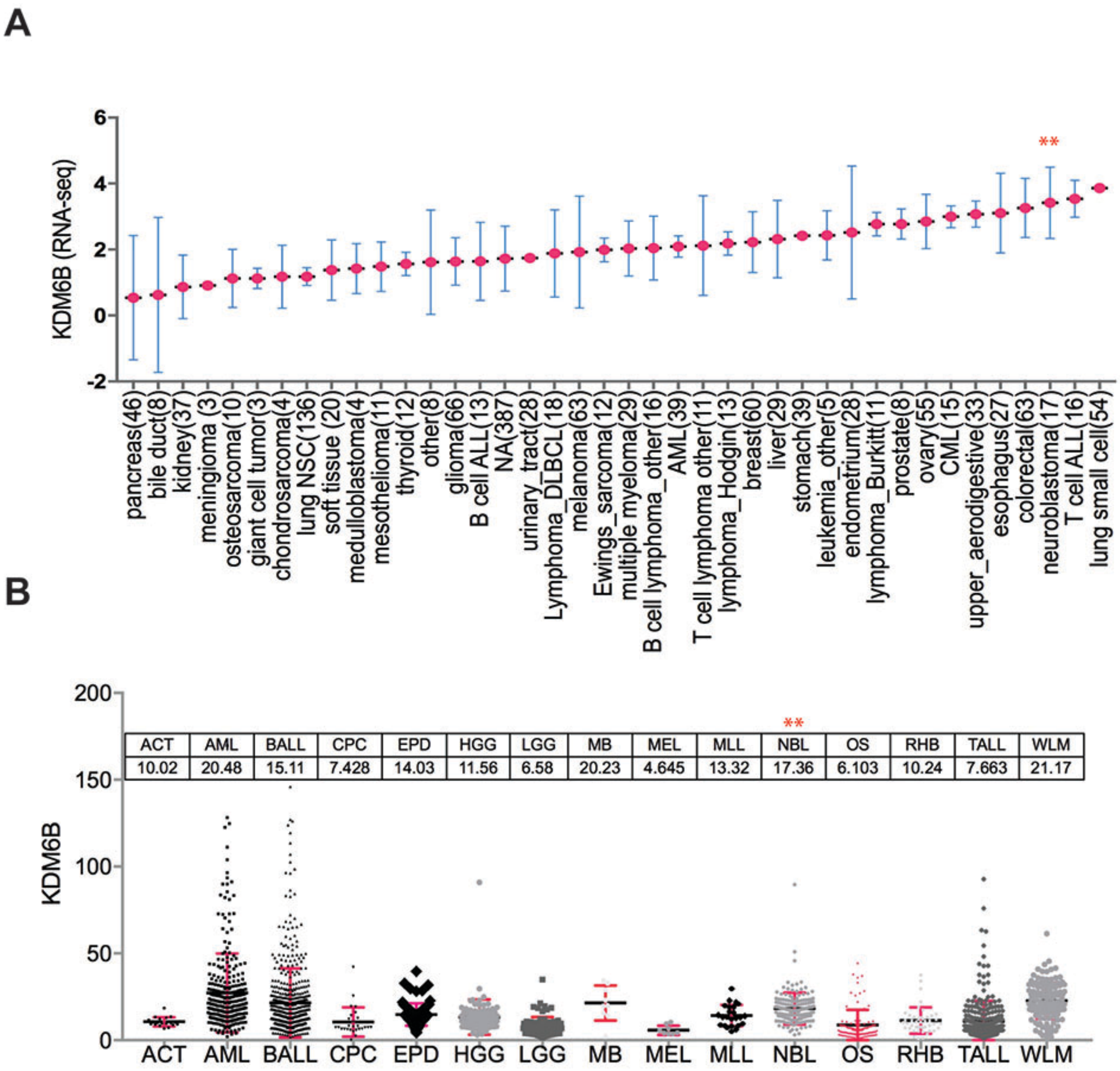


**Supplementary Figure S1. KDM6B is highly expressed in human neuroblastoma.**

(**A**) The expression of KDM6B from different cancer lineages. The RNA-seq data was downloaded from https://portals.broadinstitute.org/ccle. Y-axis represents the Fragment Per Kilobase of transcript per Million (FPKM) mapped reads.

(**B**) The expression of KDM6B from different cancer lineages. The RNA-seq data was downloaded from https://pecan.stjude.cloud. Y-axis represents the Fragment Per Kilobase of transcript per Million (FPKM) mapped reads.

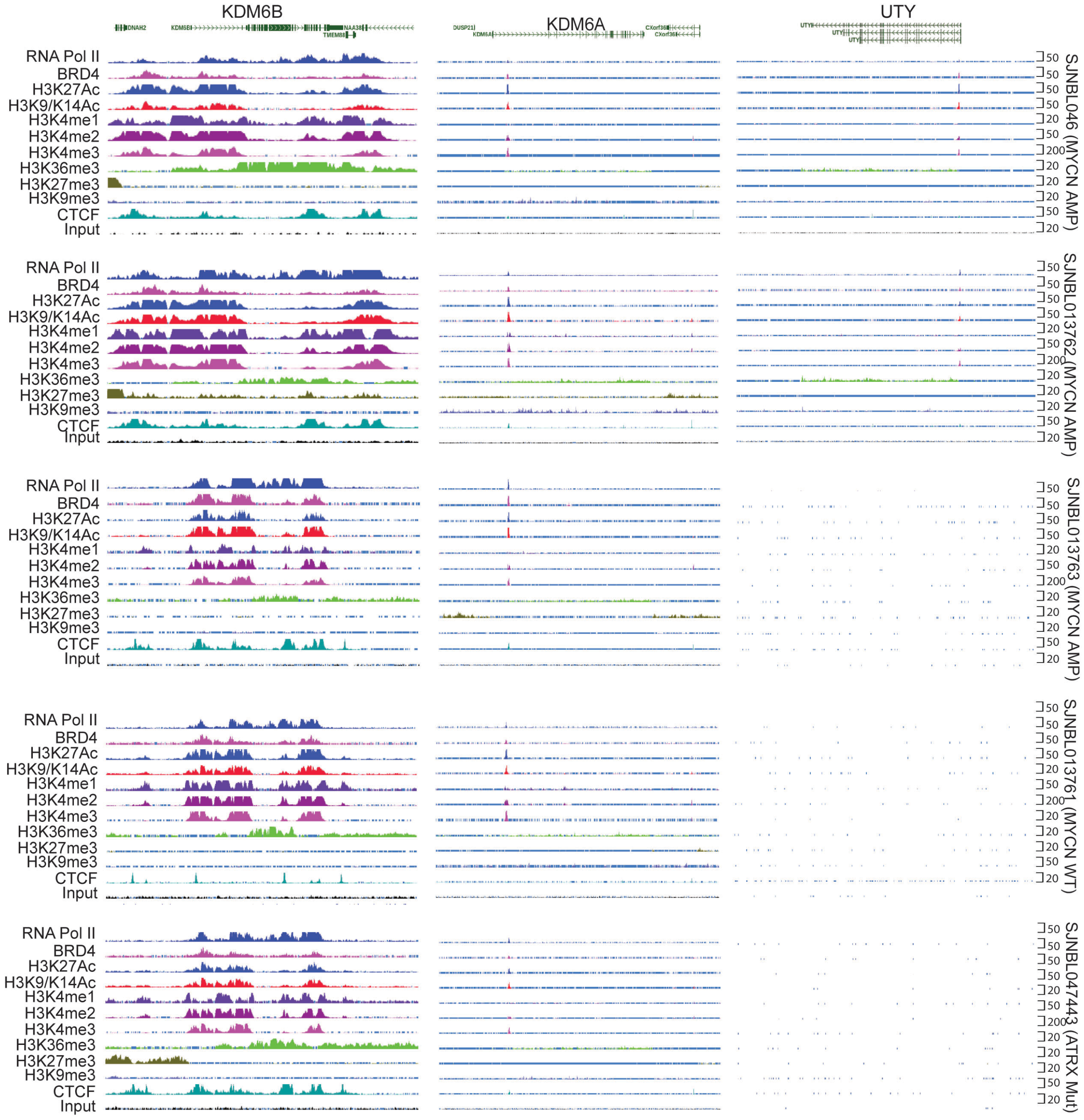


**Supplementary Figure S2. Epigenetic landscape of KDM6A, KDM6B and UTY**

The epigenetic landscapes consisting of histone marks and transcription factor binding distinguish KDM6B from KDM6A and UTY in primary neuroblastoma tissues with MYCN amplification (SJNBL046, SKNBL013762, SJNBL013763), without MYCN amplification (SJNBL013761), or with ATRX mutation (SJNBL047443).

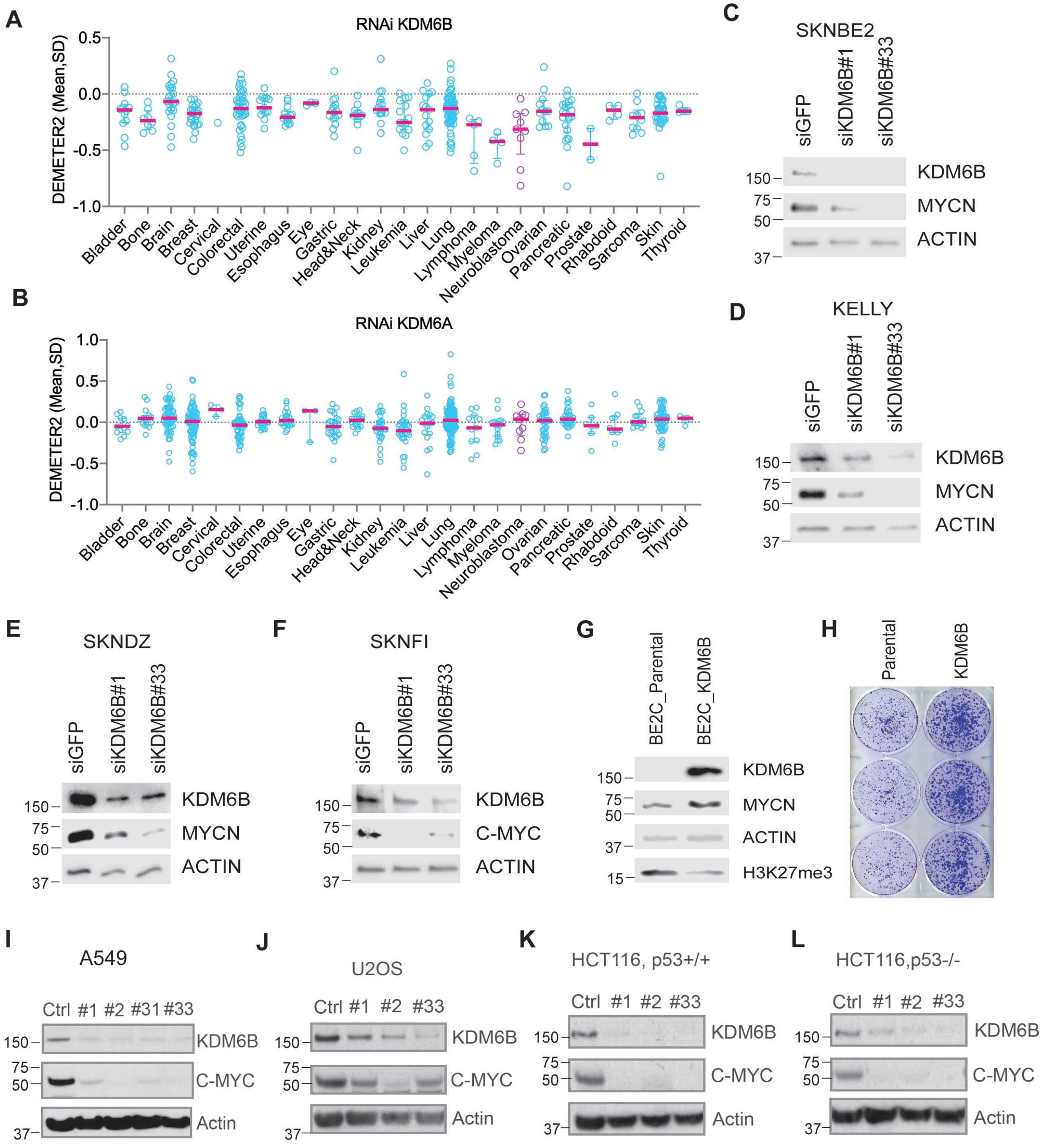


**Supplementary Figure S3. KDM6B regulates MYC expression.**

(**A-D**) The DEMETER2 score of an RNAi screen across 25 cancer lineages. DEMETER2 is a model that scores gene dependencies in 712 cancer cell lines that have been evaluated in three different large-scale pooled RNAi screens (DepMap.org).

(**C-F**) Western blot analysis with indicated antibodies to assess MYCN or C-MYC expression after 3-day transfection of 2 different siRNAs to knockdown KDM6B in neuroblastoma cell lines.

(**G**) Western blot analysis with indicated antibodies after BE2C cells are transduced with MSCV-KDM6B.

(**H**) BE2C cells and KDM6B overexpressing cells were seeded at equal numbers in a 6-well plate. 5 days later, cells were stained with crystal violet.

(**I-L**) Western blot analysis with indicated antibodies to assess MYCN or C-MYC expression after 3-day transfection of 2 different siRNA to knockdown KDM6B in lung adenocarcinoma A549 (I), osteosarcoma U2OS (J), colorectal HCT116 (K) and p53 null isogenic HCT116 cells (L).

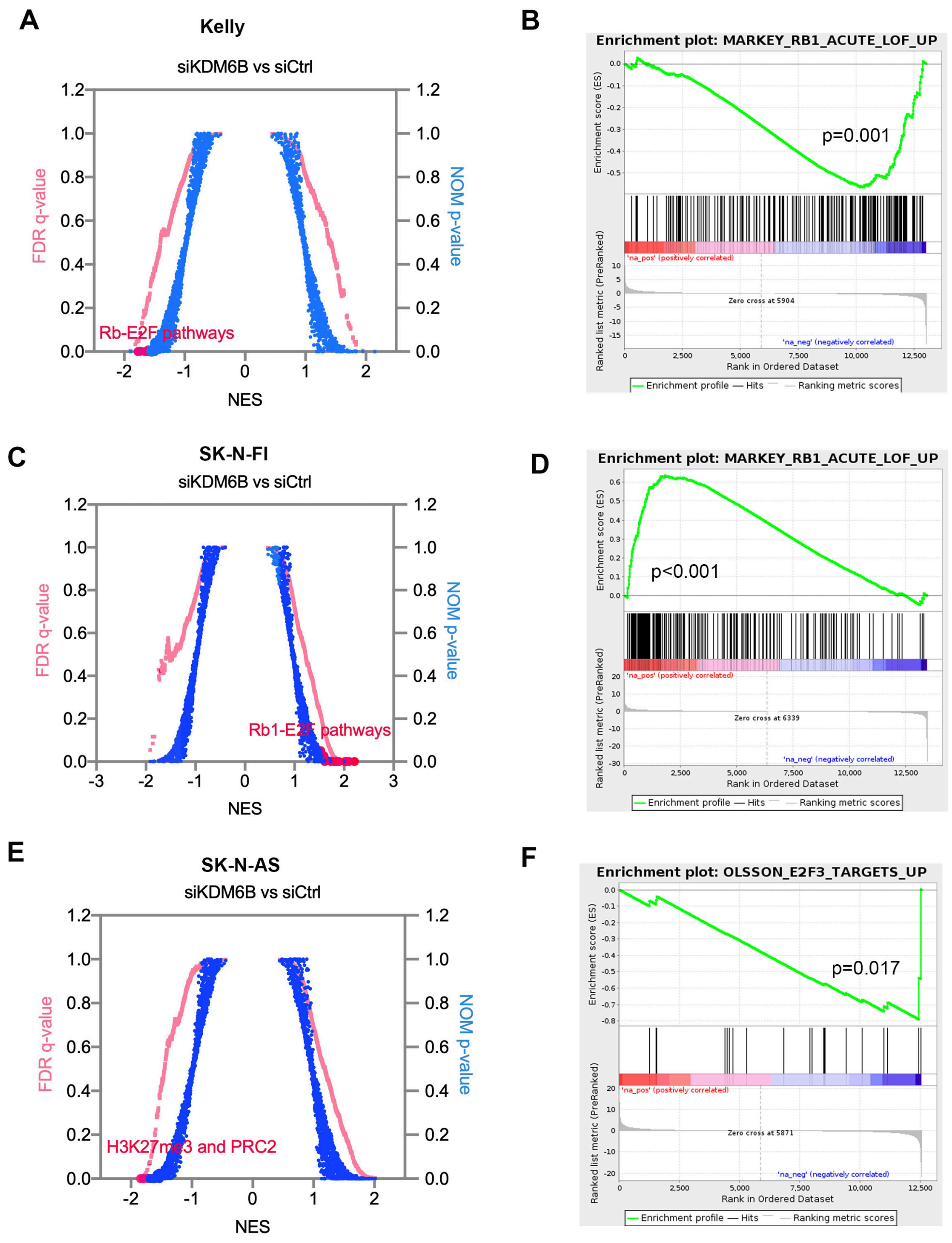


**Supplementary Figure S4. GSEA analyses for KELLY, SK-N-FI and SK-N-AS cells.**

(**A, C, E**) Quantitative comparison of all chemical and genetic perturbation gene sets from the MSigDB by gene set enrichment analysis (GSEA) for reduced (left) and increased (right) expression of global genes caused by KDM6B knockdown in KELLY (A), SK-N-FI (B) and SK-N-AS (E) cell lines. Data are presented as a scatterplot of normalized p value/false discovery q value vs normalized enrichment score (NES) for each evaluated gene set. The gene sets circled in red color indicate cell cycle, pRB-E2F and MYC pathway gene sets. (**B, D, F**) Representative GSEA samples for Kelly (B), SK-N-FI (D) and SK-N-AS (F).

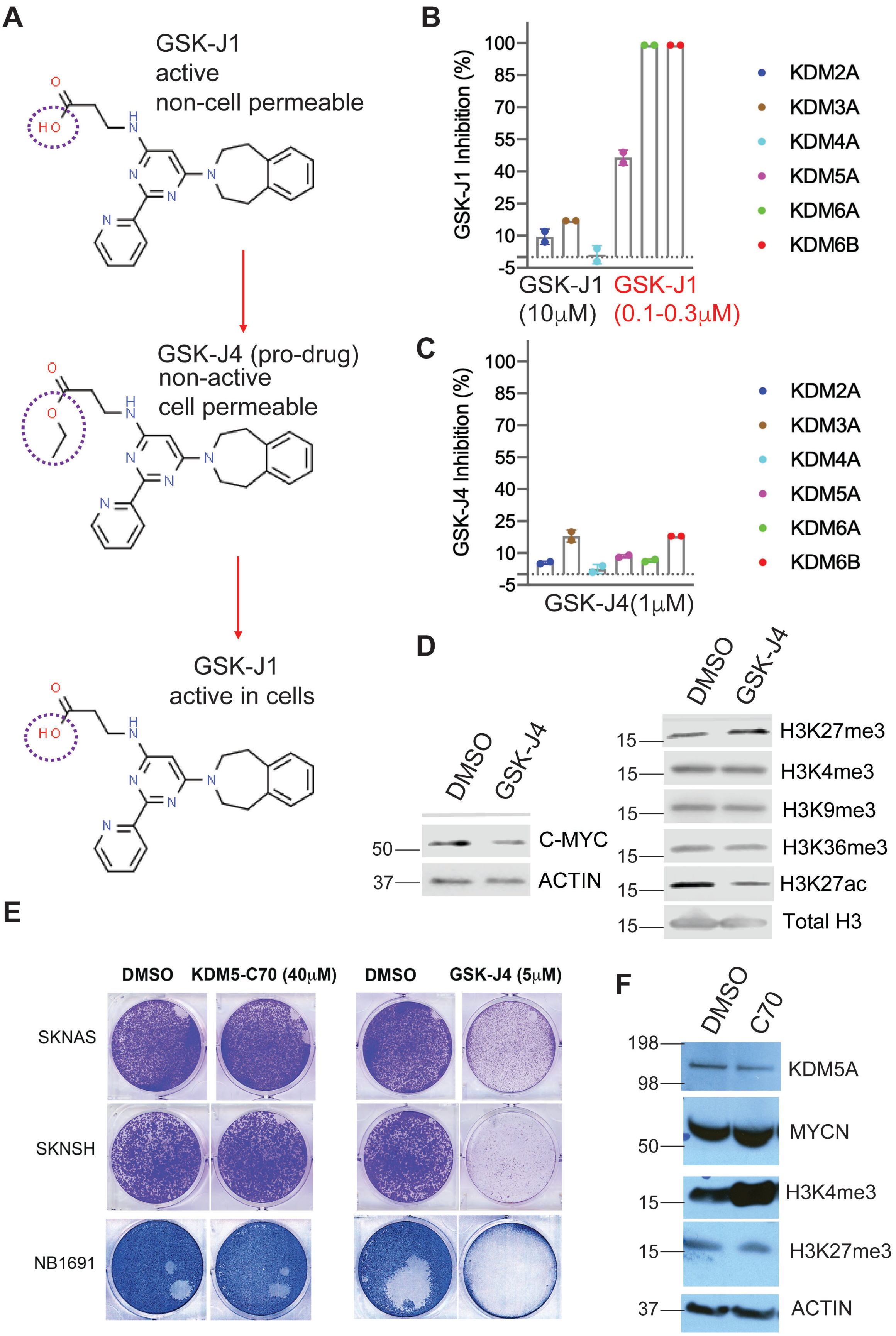
A
GSK-J1
active
non-cell permeable
GSK-J4 (pro-drug)
non-active
cell permeable
GSK-J1
active in cells
B
GSK-J1 Inhibition (%)
GSK-J1 (10μM)
GSK-J1 (0.1-0.3μM)
KDM2A
KDM3A
KDM4A
KDM5A
KDM6A
KDM6B
C
GSK-J4 Inhibition (%)
GSK-J4(1μM)
D
DMSO
GSK-J4
C-MYC
ACTIN
H3K27me3
H3K4me3
H3K9me3
H3K36me3
H3K27ac
Total H3
E
DMSO
KDM5-C70 (40μM)
GSK-J4 (5μM)
SKNAS
SKNSH
NB1691
F
C70
KDM5A
MYCN
H3K4me3
H3K27me3
ACTIN

**Supplementary Figure 5. Profiling of GSK-J1 and GSK-J4 against KDMs.**

**(A)** GSK-J1, as a selective KDM6 inhibitor, is unable to penetrate cells. To make a cell-permeable inhibitor, GSK-J1 was modified by adding an ester (named as GSK-J4) specifically to enable efficient intracellular delivery of the compound. Once entering cells, GSK-J4 converts to GSK-J1 for inhibiting KDM6.

**(B, C)** AlphasLISA applied to tests the inhibitory activity of GSK-J1 and GSK-J4 against purified KDMs in vitro, to validate the GSK-J1 selectivity against KDMs. 0.1 μM of GSK-J1 for KDM6A and KDM5A, and 0.3 μM of GSK-J1 for KDM6B.

(**D**) SK-N-AS cells were treated with GSK-J4 at 2.5μm for 48 hours. Cell lysates were subject to immunoblotting with indicated antibodies.

(**E**) SK-N-AS, SK-N-SH and NB-1691 cells were treated with 40μM of KDM5-C70 or 5μM of GSK-J4 for 10 days, followed by crystal violet staining.

(F) NB-1691 cells treated with 40μM of KDM5-C70 for 72 hours were subject to immunoblotting with indicated antibodies.

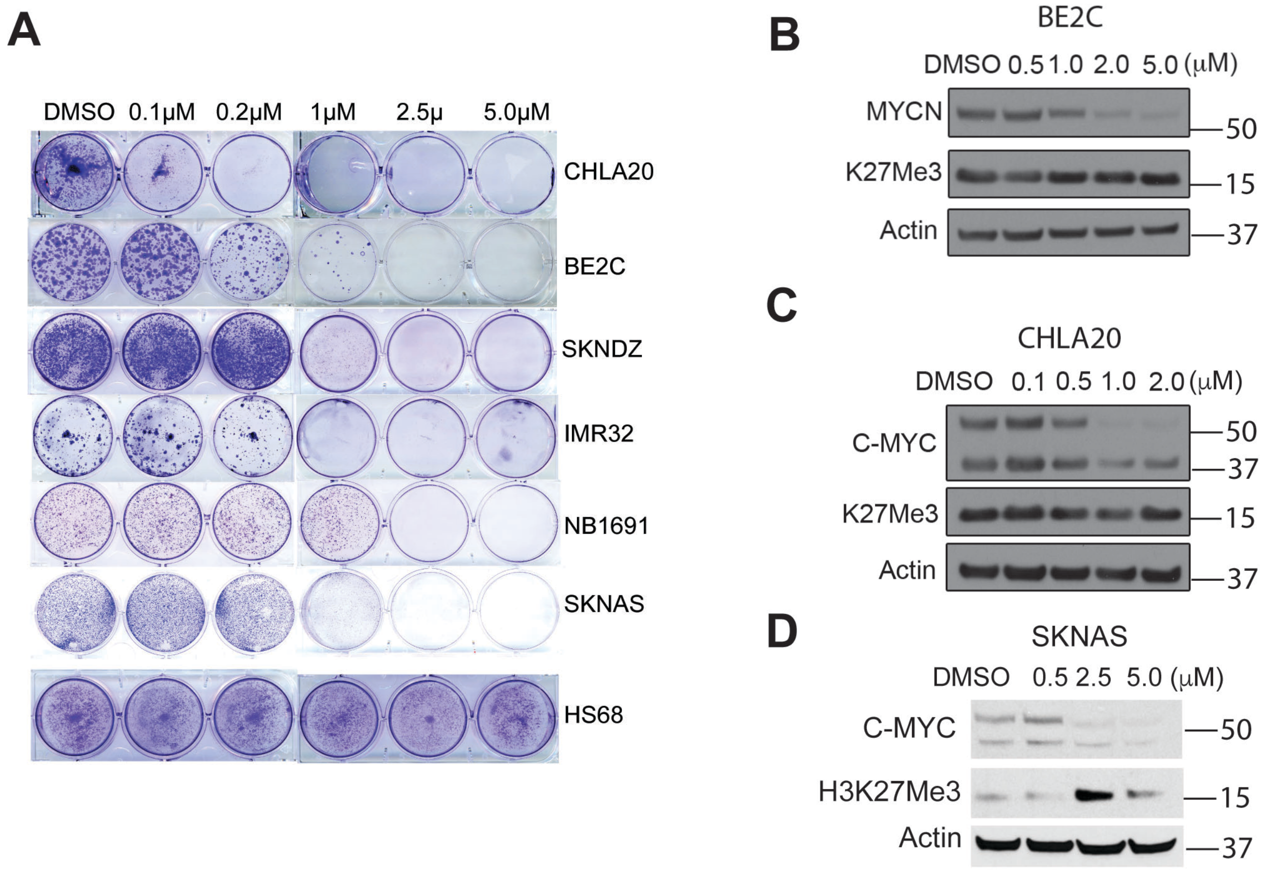


**Supplementary Figure S6. GSK-J4 targets MYC expression.**

(**A**) Crystal violet staining of colonies after neuroblastoma cell lines and human normal fibroblast HS68 cells were treated with different concentrations of GSK-J4 for 7 days.

(**B-D**) Western blot analysis with indicated antibodies to assess MYCN or C-MYC and H3K27me3 expression after 48-hour treatment with GSK-J4 of BE2C (B), CHLA20 (C) and SKNAS cells (D).

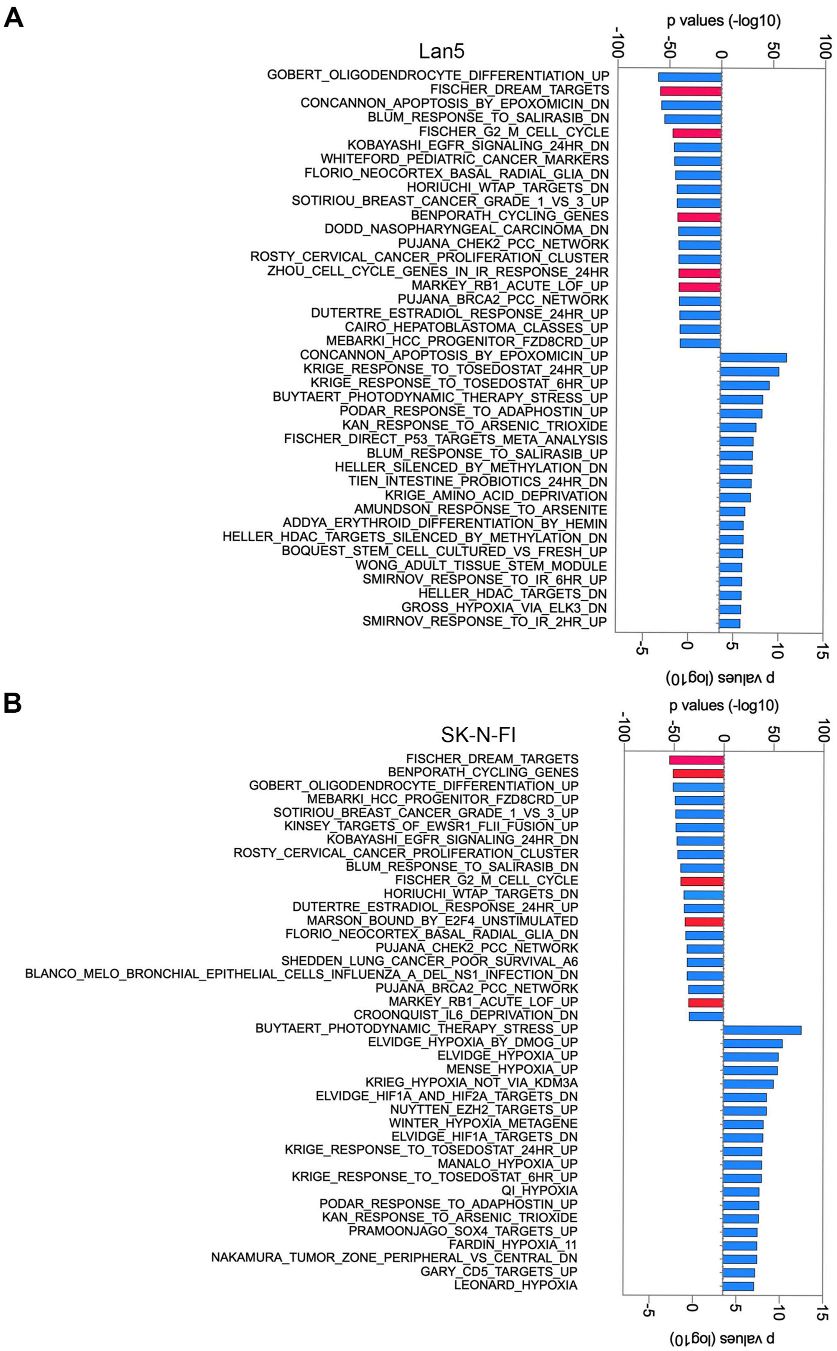


**Supplementary Figure S7. GSK-J4 inhibits E2F pathways.**

The significant differentially expressed genes (downregulated and upregulated) induced by GSK-J4 in LAN5 and SK-N-FI cells[41] were analyzed using GSEA online program (https://www.gsea-msigdb.org/gsea/msigdb/annotate.jsp) for pathway enrichment in database CGP: chemical and genetic perturbations. Red color indicates the gene sets of Rb1-E2F and cell cycle.

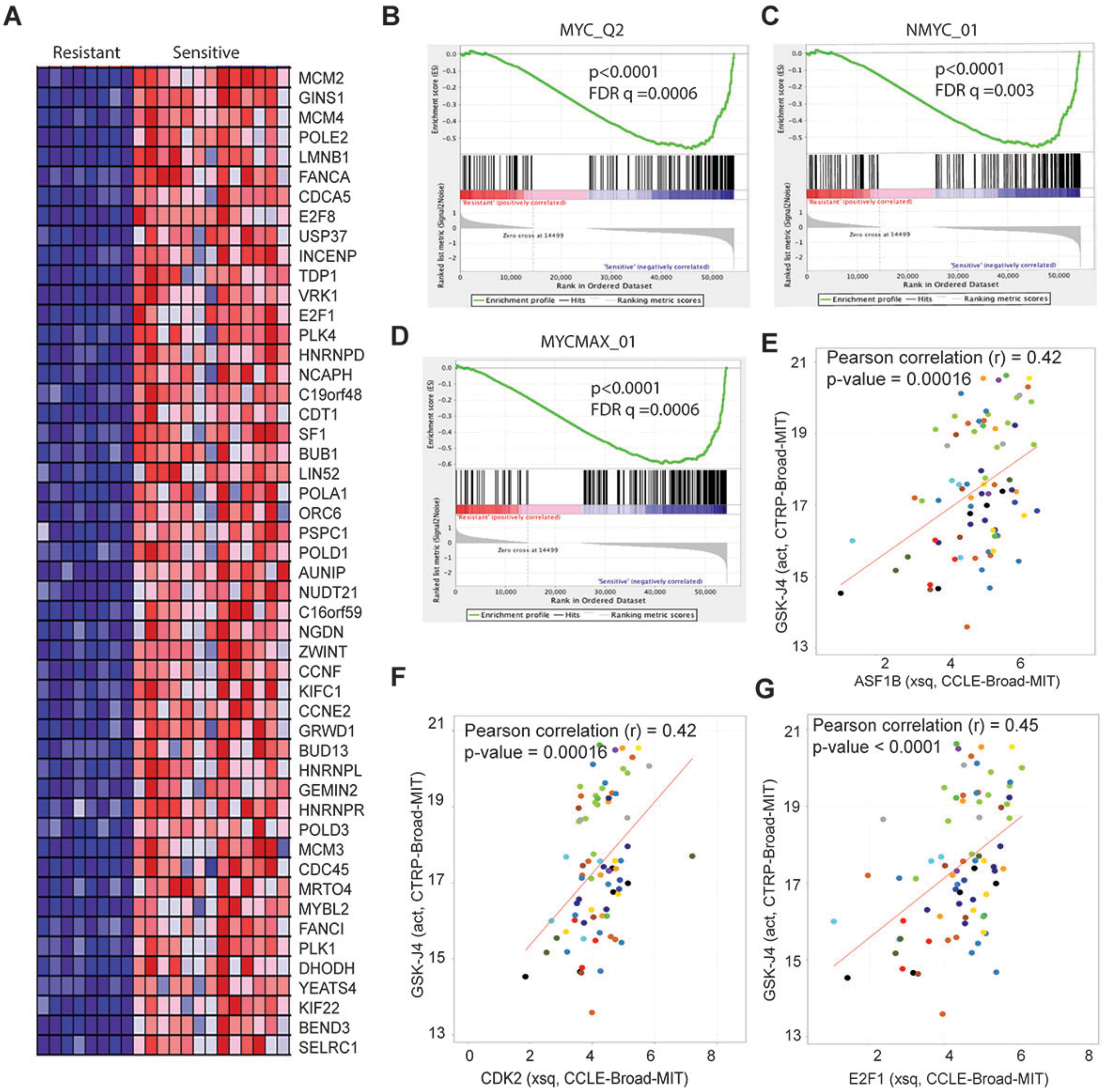


**Supplementary Figure S8. E2F signature is associated with GSK-J4 sensitivity.**

**(A)** The GSEA heatmap shows the top 50 genes highly expressed in GSK-J4 sensitive (group A) vs resistant (group B), after analysis of data which was extracted from The Cancer Therapeutics Response Portal (CTRP).

(**B-D**) GSEA show that genes highly expressed in GSK-J4 sensitive (group A) cells are enriched with MYC gene sets.

(**E-G**) The Pearson correlation of GSK-J4 with expression of E2F target genes, ASF1B(**E**), CDK2 (**F**), E2F1 (**G**), analyzed by using the online program https://discover.nci.nih.gov/cellminercdb/.

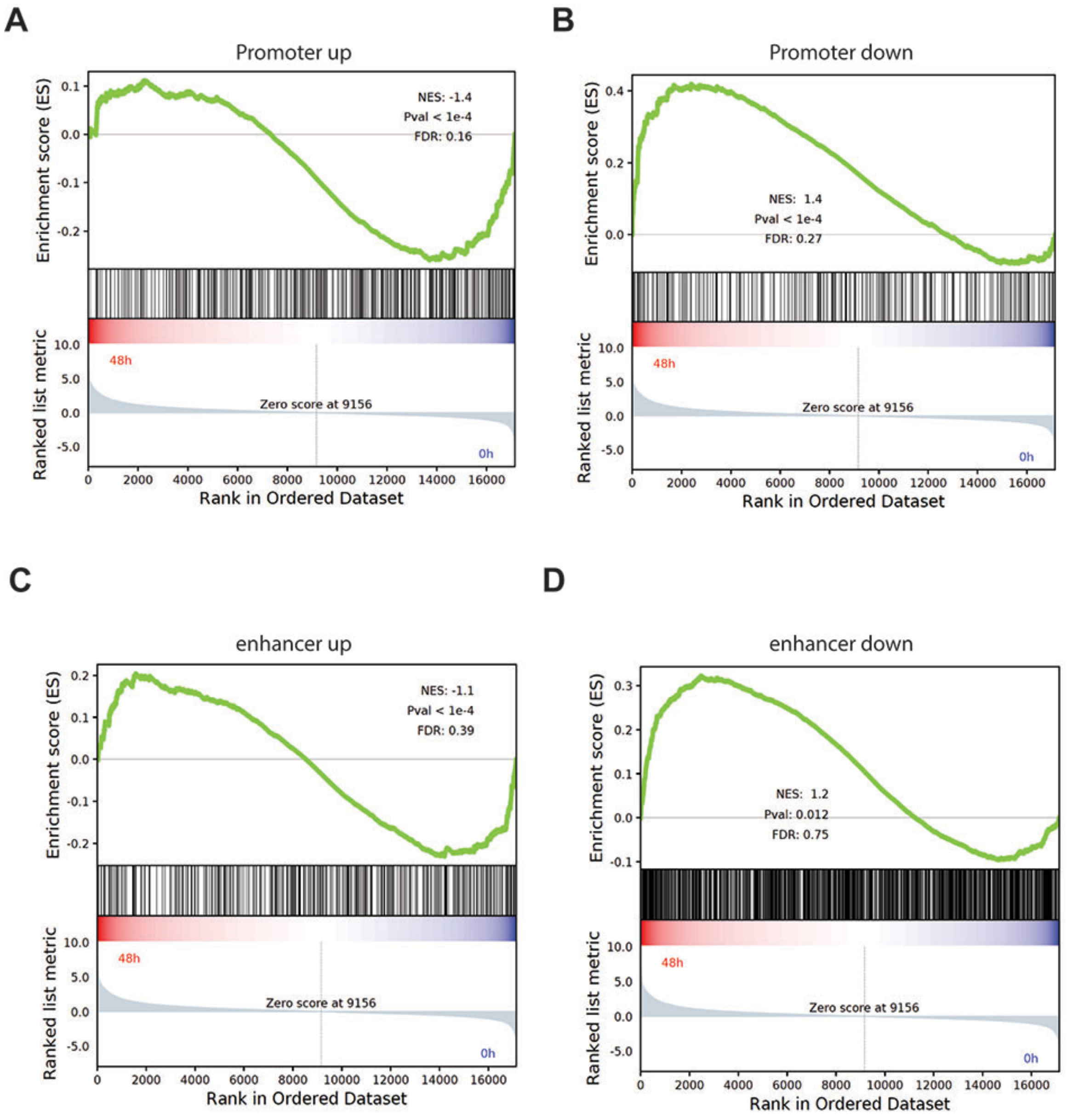


**Supplementary Figure S9.** GSEA analysis of differentially altered chromatin accessibility regions against the gene expression data based on RNA-seq at 48h treatment by GSK-J4.

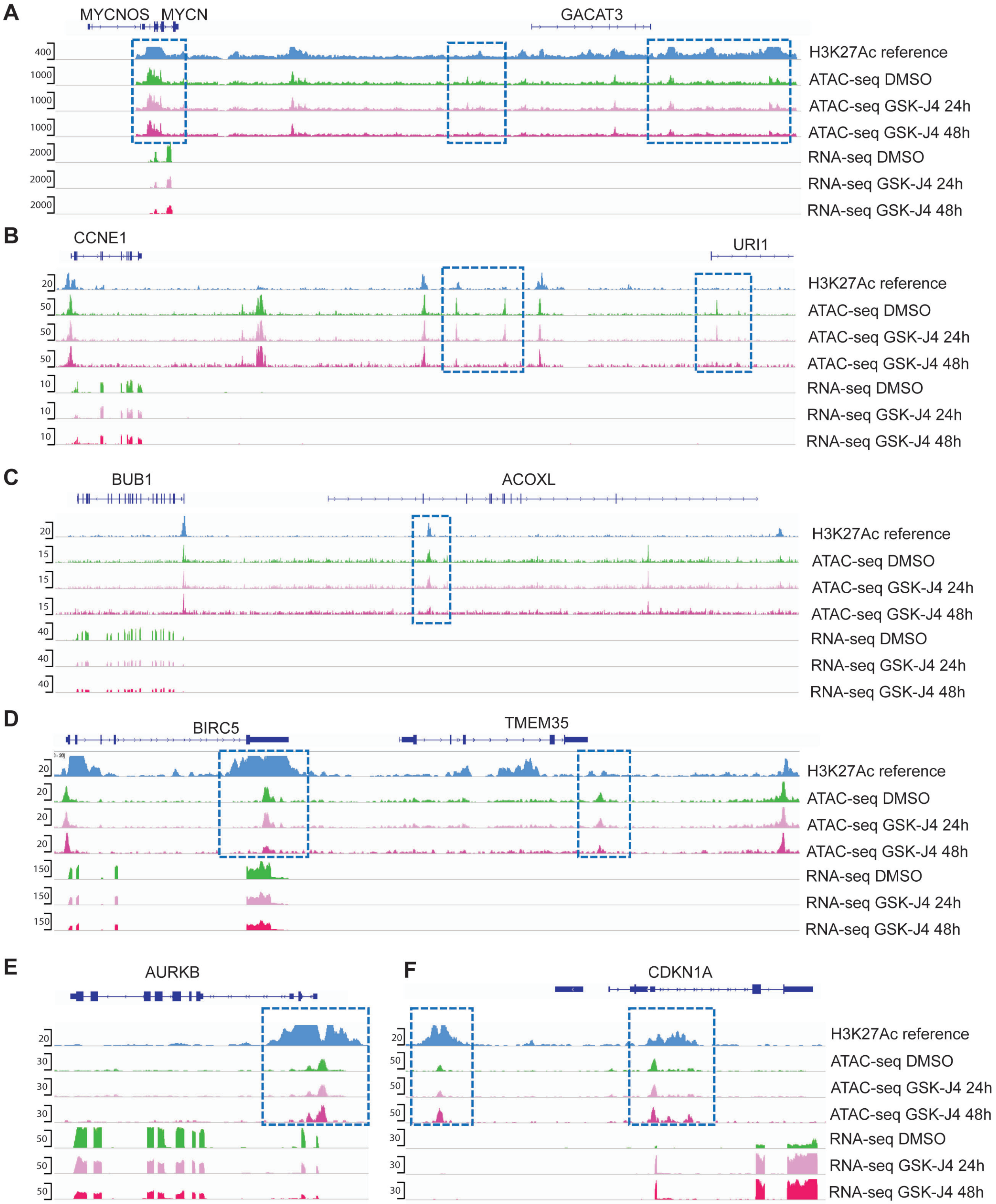


**Supplementary Figure S10.** The ATAC-seq analysis of *MYCN* and cell cycle genes (*CCNE1*, *BUB1*, *BIRC5*, *AURKB*, *CDKN1A*) shows the alterations of DNA accessibility at their promoter and distal enhancer regions by GSK-J4 treatment for 24h and 48h. RNA-seq data analysis shows the RNA reads effected by GSK-J4 treatment for 24h and 48h.

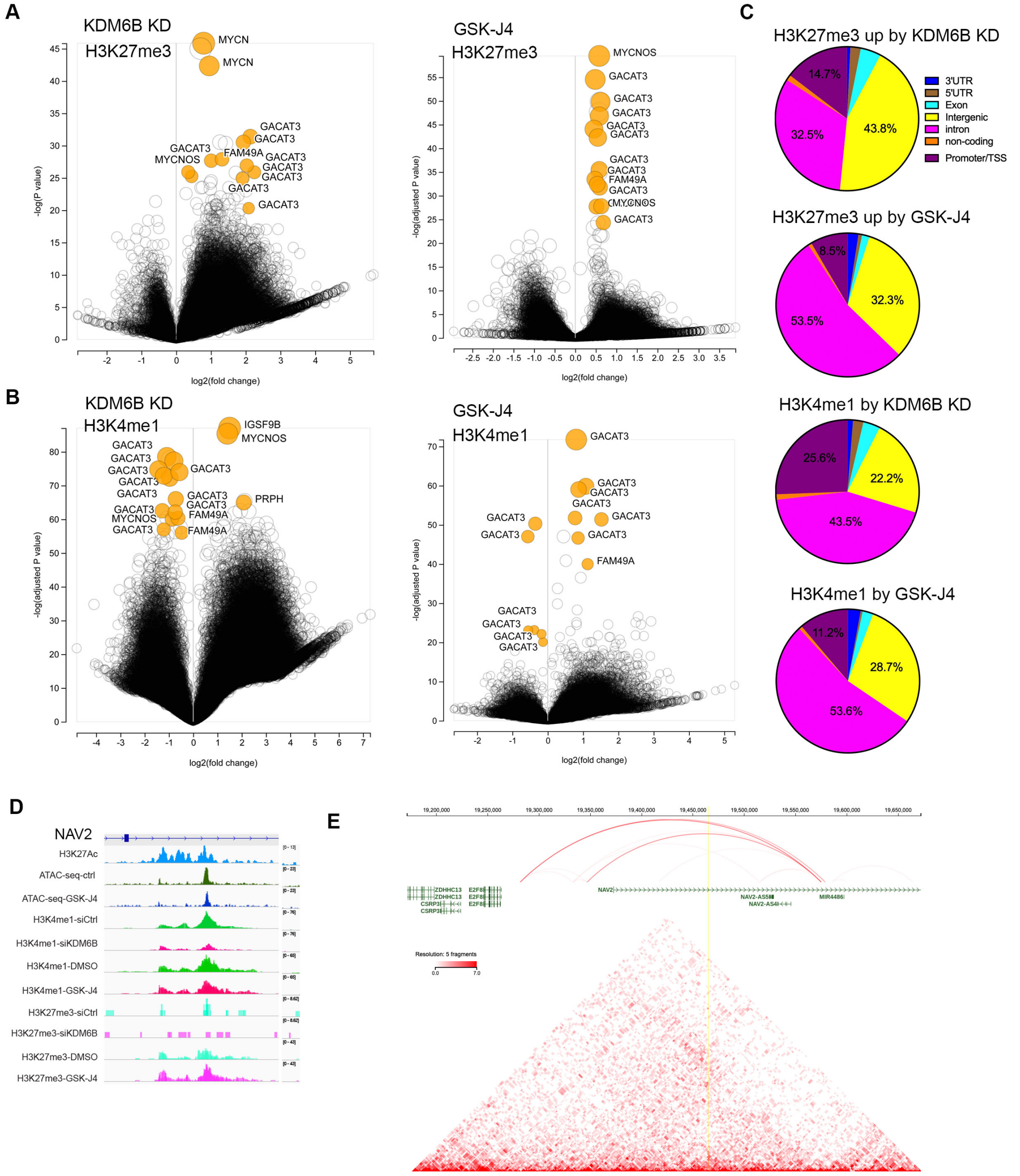

**Supplementary Figure S11. Impact of epigenetic marks by KDM6B inhibition. (A, B)** Volcano plot showing differential peaks of H3K27me3 (A) and H3K4me1 (B) induced by KDM6B knockdown and GSK-J4 treatment of BE2C cells. The yellow color highlights the super-enhancer region of *MYCN*, GACAT3 locus. (**C**) Pie plot showing the percentage of H3K27me3 upregulated by KDM6B inhibition and global H3K4me1 at distinct genomic regions. (**D**) Epigenetic snapshot of *E2F8* regulatory region, *NAV2* locus, by IGV. (**E**) The annotation of Topologically associated domain at *E2F8* and *NAV2* loci in BE2C cells (data extracted from St Jude Cloud program https://www.stjude.cloud/).

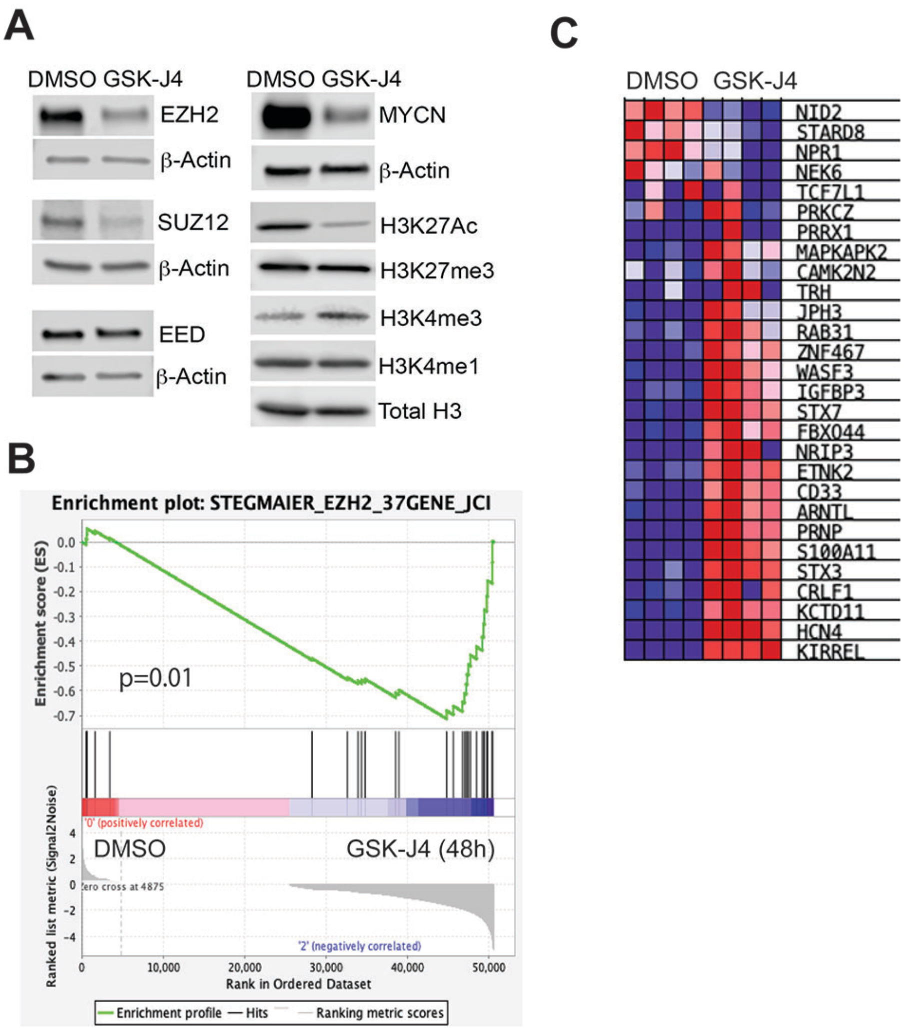


**Supplementary Figure S12. GSK-J4 downregulates the expression of PRC2 complex.**

(**A**) Western blot analysis with indicated antibodies to assess PRC2 complex, MYCN and histone modification mark expression after 48-hour treatment with 2.5µM of GSK-J4 of BE2C

(**B**) GSEA analysis of EZH2 signature against the gene expression data based on RNA-seq at 48h treatment by GSK-J4.

(**C**) Heatmap of the 37-gene signature from Figure B.

CHLA20 (C) and SKNAS cells (D) RNA-seq data shows the read peaks of each exon of SUZ12 and EED in BE2C cells treated with 2.5 µM of GSK-J4 for 24 and 48h.

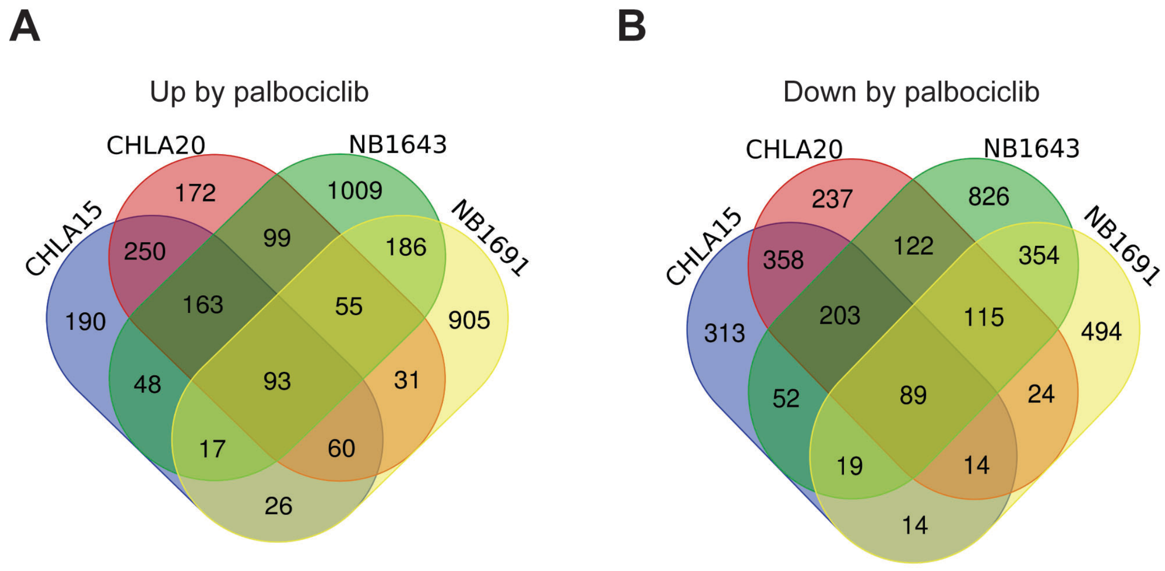


**Supplementary Figure S13. Common genes done regulated and upregulated by palbociclib.** 4 neuroblastoma cells lines were treated with palbociclib for RNA-seq. The differentially expressed genes in each cell line were analyzed by using Venn program to find the commonly upregulated (A) or downregulated genes (B)

A

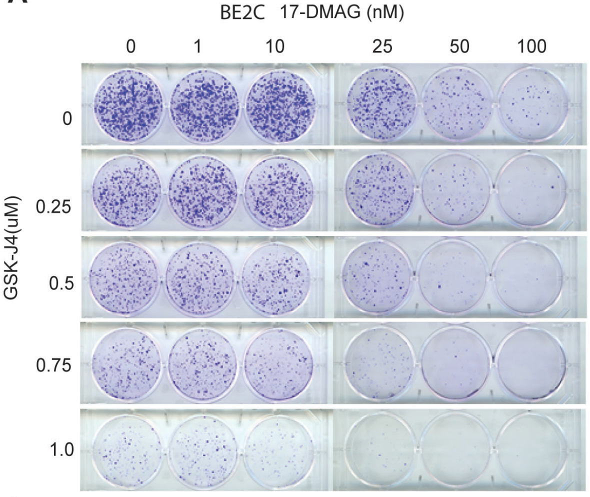


C

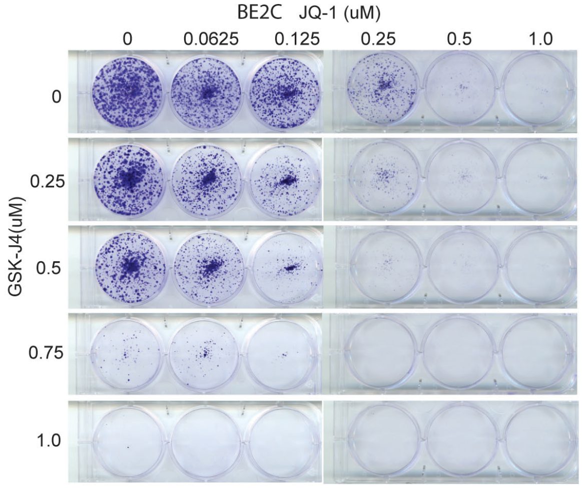


B

BE2C 17-DMAG (nM)

Percent over bliss score

| GSK-J4(uM) | 0 | 1 | 10 | 25 | 50 | 100 |
|---|---|---|---|---|---|---|
| 0 | | | | | | |
| 0.25 | | -0.1 | -1.5 | 1.5 | 3.1 | 7.3 |
| 0.5 | | 7.9 | 0.5 | 5.4 | 12.9 | 12.0 |
| 1 | | 1.7 | 9.7 | 9.7 | 6.8 | 4.8 |

D

BE2C JQ-1(uM)

Percent over bliss score

| GSK-J4(uM) | 0 | 0.0625 | 0.125 | 0.25 | 0.5 | 1 |
|---|---|---|---|---|---|---|
| 0 | | | | | | |
| 0.25 | | 3.5 | 3.5 | 4.7 | 5.7 | 6.1 |
| 0.5 | | 5.3 | 6.5 | 9.0 | 8.2 | 10.2 |
| 1 | | -6.6 | -2.6 | -2.0 | 1.0 | 7.7 |

E

SK-N-AS Palbocicilb (uM)

| GSK-J4(uM) | 0 | 0.25 | 0.5 | 1 | 2 | 4 |
|---|---|---|---|---|---|---|
| 0 | | | | | | |
| 0.5 | | 2.8 | 0.7 | 0.5 | -7.4 | -1.3 |
| 1 | | 1.9 | 0.1 | -2.8 | -2.3 | 2.7 |
| 2.5 | | -3.6 | -2.3 | -3.3 | 2.8 | 3.9 |

F

SK-N-AS 17-DMAG (nM)

| GSK-J4(uM) | 0 | 1 | 10 | 25 | 50 | 100 |
|---|---|---|---|---|---|---|
| 0 | | | | | | |
| 0.5 | | 5.2 | -3.2 | -6.4 | 2.3 | 2.0 |
| 1 | | -3.5 | -6.2 | -7.3 | 2.9 | 3.1 |
| 2.5 | | -2.8 | -7.9 | -4.7 | 1.3 | 1.9 |

G

SK-N-AS JQ-1(uM)

| GSK-J4(uM) | 0 | 0.0625 | 0.125 | 0.25 | 0.5 | 1 |
|---|---|---|---|---|---|---|
| 0 | | | | | | |
| 0.5 | | -2.8 | -1.4 | 0.8 | -1.2 | -3.8 |
| 1 | | -1.1 | -3.4 | -1.4 | -7.0 | 2.1 |
| 2.5 | | 0.6 | 2.1 | -1.5 | 3.6 | 4.3 |

**Supplementary Figure S14. KDM6 inhibition is synergistic with non-CDK4/6 inhibitors**

**(A)** BE2C cells were seeded with low numbers in 6-well plate and treated with different concentrations of GSK-J4 or/and 17-DMAG for 7 days. The cell colonies were stained with crystal violet.

(**B**) Bliss index for combination of GSK-J4 and 17-DMAG in BE2C cells. Positive scores indicate synergy, negative scores indicate antagonism.

(**C**) BE2C cells were seeded with low numbers in 6-well plate and treated with different concentrations of GSK-J4 or/and JQ-1 for 7 days. The cell colonies were stained with crystal violet.

(**D**) Bliss index for combination of GSK-J4 and JQ-1 in BE2C cells. Positive scores indicate synergy, negative scores indicate antagonism.

(**E**) Bliss index for combination of GSK-J4 and palbociclib in SK-N-AS cells. Positive scores indicate synergy, negative scores indicate antagonism.

(**F**) Bliss index for combination of GSK-J4 and 17-DMAG in SK-N-AS cells. Positive scores indicate synergy, negative scores indicate antagonism.

(**G**) Bliss index for combination of GSK-J4 and JQ-1 in SK-N-AS cells. Positive scores indicate synergy, negative scores indicate antagonism.

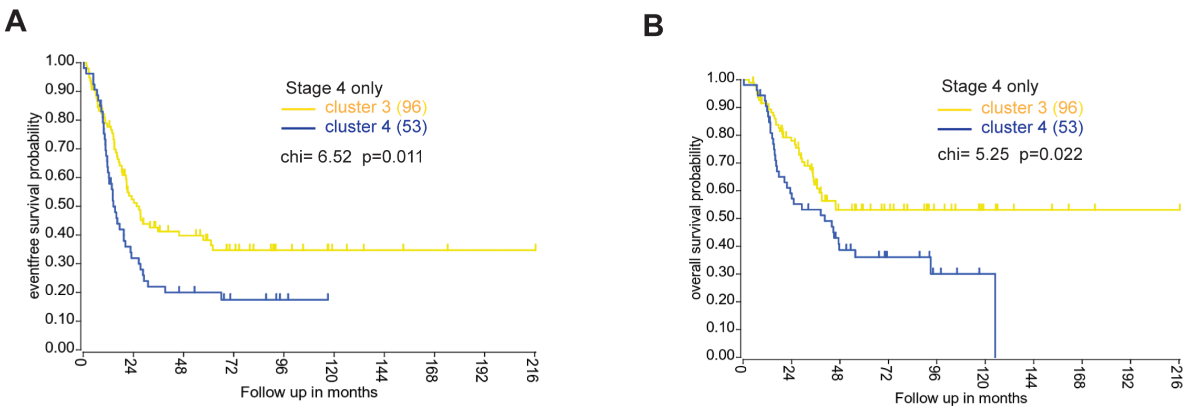


**Supplementary Figure 15**. Kaplan-Meier curve using Log-Rank method shows the event-free survival and overall survival of clusters 3 and 4 with stage 4 disease.

**Supplemental Table S1. Cell lines related to Figure 3B.**

| | |
|---|---|
| Resistant cells | KYSE30_OESOPHAGUS |
| | TUHR14TKB_KIDNEY |
| | BICR56_UPPER_AERODIGESTIVE_TRACT |
| | HS940T_FIBROBLAST |
| | HS688AT_FIBROBLAST |
| | KMRC3_KIDNEY |
| | KG1C_CENTRAL_NERVOUS_SYSTEM |
| | HS822T_FIBROBLAST |
| Sensitive cells | KASUMI2_HAEMATOPOIETIC_AND_LYMPHOID_TISSUE |
| | HGC27_STOMACH |
| | KYM1_SOFT_TISSUE |
| | COV434_OVARY |
| | DMS53_LUNG |
| | A204_SOFT_TISSUE |
| | KMBC2_URINARY_TRACT |
| | T84_LARGE_INTESTINE |
| | MDAMB453_BREAST |
| | DMS114_LUNG |
| | KE39_STOMACH |
| | SKNBE2_AUTONOMIC_GANGLIA |
| | SNU216_STOMACH |

**Supplemental Table S2. Transcription factor motif (TFT) for group A cells in Figure 3B**

| | Gene sets | NES | NES | NOM p-val | FDR q-val |
|---|---|---|---|---|---|
| 1 | E2F_Q3 | -0.73 | -2.27 | 0 | 0 |
| 2 | E2F_Q4_01 | -0.74 | -2.24 | 0 | 0 |
| 3 | E2F_Q6_01 | -0.71 | -2.24 | 0 | 0 |
| 4 | E2F1_Q6 | -0.77 | -2.23 | 0 | 0 |
| 5 | E2F1_Q3 | -0.76 | -2.23 | 0 | 0 |
| 6 | E2F1_Q4_01 | -0.72 | -2.23 | 0 | 0 |
| 7 | E2F1_Q6_01 | -0.76 | -2.22 | 0 | 0 |
| 8 | E2F_Q3_01 | -0.72 | -2.22 | 0 | 0 |
| 9 | SGCGSSAAA_E2F1DP2_01 | -0.78 | -2.21 | 0 | 0 |
| 10 | E2F1DP1_01 | -0.75 | -2.21 | 0 | 0 |
| 11 | E2F1DP2_01 | -0.75 | -2.21 | 0 | 0 |
| 12 | E2F4DP2_01 | -0.75 | -2.21 | 0 | 0 |
| 13 | E2F4DP1_01 | -0.75 | -2.21 | 0 | 0 |
| 14 | E2F_03 | -0.72 | -2.21 | 0 | 0 |
| 15 | E2F_Q6 | -0.76 | -2.21 | 0 | 0 |
| 16 | E2F_02 | -0.75 | -2.2 | 0 | 0 |
| 17 | E2F_Q4 | -0.75 | -2.19 | 0 | 0 |
| 18 | E2F1DP1RB_01 | -0.73 | -2.19 | 0 | 0 |
| 19 | KTGGYRSGAA_UNKNOWN | -0.71 | -2.17 | 0 | 0 |
| 20 | E2F_01 | -0.76 | -2.15 | 0 | 0 |
| 21 | E2F1_Q4 | -0.66 | -2.11 | 0 | 0 |
